# There and Back to the Present: An Evolutionary Tale on Biological Diversity

**DOI:** 10.1101/2021.12.11.472171

**Authors:** Leandro Duarte, Gabriel Nakamura, Vanderlei Debastiani, Renan Maestri, Maria João Ramos Pereira, Marcus Cianciaruso, José Alexandre F. Diniz-Filho

**Affiliations:** Programa de Pós-graduação em Ecologia, Instituto de Biociências, Universidade Federal do Rio Grande do Sul, Av. Bento Gonçalves 9500 CP 15007 Porto Alegre, 91501-970, Brazil; Programa de Pós-graduação em Ecologia e Evolução, Instituto de Ciências Biológicas, Universidade Federal de Goiás. CP 131, 74690-900, Goiânia, GO, Brazil

**Keywords:** Ornstein-Uhlenbeck model, adaptive rate, Approximate Bayesian Computation, eco-evolutionary dynamics

## Abstract

Ecologists often agree on the importance of macroevolution for niche-mediated distribution of biological diversity along environmental gradients. Yet, macroevolutionary diversification and dispersal in time and space generate uneven geographic distribution of phylogenetic pools, which affects the imprint let by macroevolution on local species pools. In this article we introduce an individual-based simulation approach coupled to Approximate Bayesian Computation (ABC) that allows to parameterize the adaptation rate of species’ niche positions along the evolution of a monophyletic lineage, and the intensity of dispersal limitation, associated with the distribution of biological diversity between assemblages potentially connected by dispersal (metacommunity). The analytical tool was implemented in an R package called *mcfly*. We evaluated the statistical performance of the analytical framework using simulated datasets, which confirmed the suitability of the analysis to estimate adaptation rate and dispersal limitation parameters. Further, we evaluated the role played by niche evolution and dispersal limitation on species diversity distribution of Phyllostomidae bats across the Neotropics. The framework proposed here shed light on the links between niche evolution, dispersal limitation and the distribution of biological diversity, and thereby improved our understanding of evolutionary imprints on ecological patterns. Perhaps more importantly, it offers new possibilities for solving the eco-evolutionary puzzle.

## Introduction

Why some places have so many species while others have so few? This is a central question that generations of ecologists after G. E. Hutchinson have been trying to answer after his seminal article (Hutchinson 1959). Indeed, biological diversity is the most widely addressed feature of ecological assemblages; yet, its interpretation may be tricky (Ricotta 2005). The mechanisms determining the striking variation of biological diversity over space and across environmental gradients have been debated for decades (Vellend 2016). Alternative (and complementary) theories have been proposed to explain biological diversity, either based on species niches (Hutchinson 1957; Levine and HilleRisLambers 2009), neutral processes (Hubbell 2001; MacArthur and Levins 1967) or biogeographic and historical factors (Gerhold et al. 2018; MacArthur 1972; Ricklefs and Schluter 1993).

Over the last decades, community phylogenetics (Cavender-Bares et al. 2009; Mouquet et al. 2012; Webb et al. 2002) has posited that phylogenetic relationships among sympatric species may link niche evolution to biogeography and, ultimately, help understanding patterns in regional and local biological diversity patterns. Accordingly, niche divergence between closely related species may be somehow prevented along evolutionary history (Felsenstein 1985; Harvey and Pagel 1991; Nosil 2012). Such tendency of species niche to vary less than expected by chance along evolution is often interpreted as evidence of phylogenetic niche conservatism (Losos 2008; Pyron et al. 2015; Wiens and Graham 2005). If so, and whether dispersal plays only a minor role in species assembly process (which is very disputable, see Vellend 2016), phylogenetic niche conservatism would provide an explicit causal link between the phylogenetic diversity of local communities and major ecological mechanisms (environmental filtering, biotic interactions, see Webb et al. 2002). Following this rationale, some ecologists have been advocating the use of phylogenetic covariation as a proxy for niche similarity among species (Cadotte et al. 2012; Tucker et al. 2018).

Nonetheless, evidence against such ‘phylogeny-as-proxy’ approach has been accumulated over the last years, as similar phylogenetic diversity patterns may be explained by different, and often opposing, mechanisms (Gerhold et al. 2015; Godoy et al. 2014; Mayfield and Levine 2010; Swenson 2019). As controversial conclusions on the usefulness of interpreting phylogenetic diversity in the light of ecological mechanisms promoting biological diversity proliferated, further theoretical advances in community phylogenetics have decelerated over the last years (but see Cadotte et al. 2017; Tucker et al. 2018). Tracing back plausible evolutionary scenarios underlying species distribution patterns across current environmental gradients remains a main challenge for community phylogenetics (Webb et al. 2002).

Unravelling evolutionary processes underlying niche divergence might help explaining the distribution of extant species belonging to a given clade across ecological gradients (McPeek 2017; Weber et al. 2017). Nonetheless, such enterprise faces two big challenges. First, theoretical and empirical evidence often demonstrate that the inference of eco-evolutionary niche dynamics based on static estimates of phylogenetic signal in species traits is not unequivocal (Münkemüller et al. 2015; Revell et al. 2008). This is because strong phylogenetic signal in species traits can emerge from processes other than niche conservatism, including neutral evolution (Cooper et al. 2010; Gould and Lewontin 1979; Harvey and Pagel 1991; Nosil 2012). Indeed, under high adaptive rate pulling species niches to an optimum value (which might be also interpreted as niche conservatism), phylogenetic signal in the niche is completely vanished (Butler and King 2004; Hansen et al. 2008). Second, ecologists often measure species traits to characterize their niches (Elton 1927; McPeek 2017), under the sound premise that species niches are mostly molded by ecological pressures determining phenotypic patterns of species (Ackerly 2003; Hansen and Martins 1996). In this sense, trait dominance and/or complementarity across environmental gradients (Garnier et al. 2016) is often taken as evidence of niche-mediated species assembly patterns. However, species traits correlate among themselves due to their shared evolutionary history, which results in complex trait-environment relationships. Therefore, inferring which specific trait(s) explain the distribution of biological diversity across environmental gradients is not trivial (Duarte et al. 2018). Indeed, searching for an evolutionary basis of niche-based species assembly across environmental gradients is still a topic of debate among ecologists (Cadotte et al. 2017; McPeek 2017; Tucker et al. 2018; Weber et al. 2017).

If neither phylogenetic signal in niche traits nor trait-environment correlations allow to build a solid bridge between niche evolution and species assembly across environmental gradients, then what? A starting point is to consider under which circumstances we should expect a detectable imprint of evolutionary processes on biological diversity. First, such imprint only makes sense if we consider a monophyletic lineage with most branches well represented across the geographic space under analysis. Otherwise, we may be investigating phylogenetic signal in biological diversity based on species (1) whose evolution was independent for the most time lineages have been diversifying, for example, tree ferns and angiosperms (Duarte et al. 2012) or Anura and Gymnophiona (Loyola et al. 2013), or (2) whose phylogenetic relationships are too incomplete or fragmented to provide a realistic evolutionary scenario (Crisp and Cook 2005). However, even when the species pool belongs to a monophyletic, well represented lineage, endemic to the region or biome we are interested in, the imprint left by phylogeny on regional or local diversity patterns may either persist or be completely erased (Maestri and Duarte 2020). This is because species assembly, and therefore biological diversity, is an instantaneous snapshot of complex eco-evolutionary dynamics acting upon co-occurring species (Shipley 2010; Weber et al. 2017). If the adaptation rate of species from an ancestral niche position towards an optimum niche condition, mediated by short-term biotic interactions (Paine et al. 2018) and/or environmental filters (Kraft et al. 2015), was sufficiently strong to break phylogenetic signal in species niches (Hansen et al. 2008), no phylogenetic imprint on species assembly would be detected. Thus, it is not surprising that tracing back evolutionary processes underlying biological diversity patterns based on static phylogenetic signal among species may lead to disparate conclusions (Gerhold et al. 2015). Appropriate tools for the evaluation of the influence of phylogeny on biological diversity across space are necessary for this purpose.

A possible solution to solve this puzzle implies building alternative scenarios of niche evolution given a phylogenetic hypothesis to generate expected species assembly patterns. This should allow us to infer the most likely scenarios of niche evolution determining empirical species assembly patterns based on the set of expectations that most resemble the empirical data. In this article we introduce a model-based simulation approach that allows to estimate the strength of adaptation rate along the niche evolution of species belonging to a monophyletic lineage and distributed across metacommunities, defined as sets of local assemblages connected by dispersal (Leibold et al. 2004). Moreover, the new framework also allows to estimate to what extent dispersal limitation influences diversity distribution across space. We start by defining the theoretical aspects of niche- and neutral-based species assembly into local assemblages, which provide the necessary background to a further contextualization of niche evolution as a component of niche-based species assembly. Afterwards, we introduce a rejection-based Approximate Bayesian Computation (ABC) algorithm (Beaumont 2010; Csilléry et al. 2010) to estimate the strength of adaptation rate and dispersal limitation given a set of sites described by species belonging to a monophyletic clade, a phylogenetic hypothesis for these species, spatial coordinates of sites, and an environmental factor associated with species assembly patterns. We demonstrate the usefulness of the simulation approach for inferring the imprint left by niche evolution and dispersal limitation on species diversity patterns across metacommunity and biogeographical scales by evaluating the statistical performance of the ABC algorithm using simulated datasets. Further, we investigate the role played by niche evolution and dispersal limitation on species diversity distribution of Phyllostomidae bats across the Neotropics.

### Parameterizing niche and neutral processes determining species assembly

Niche theory partially explains in part the distribution of biological diversity across space. The availability of key resources and the prevalence of specific levels of environmental conditions in a site enable populations of some species to thrive, while others decrease their numbers for the same reason. Together with environmental filtering, biotic interactions modulate the probability of maintenance of viable populations of some species in the presence of others (Ackerly 2003; Hutchinson 1957; MacArthur and Levins 1967; Vellend 2016; Violle et al. 2012, but see Kraft et al. 2015).

On the other hand, neutral dynamics relies on dispersal-based species assembly to explain biological diversity. Neutral dynamics is modeled based on dispersal limitation, immigration rates, local and regional species abundances, and speciation events determining demography in local species assemblages, assuming that species are ecologically equivalent in respect to their niches (Hubbell 2001). Even though no one dispute the importance of niche-based processes to explain species assembly patterns, dispersal processes also play an important role to determine species distribution across space (Vellend 2016). Therefore, the importance of neutral processes (Bell 2000; Hubbell 2001) cannot be underestimated. In fact, such dichotomy between niche and neutral processes is only apparent (Gravel et al. 2006), since these processes represent complementary mechanisms determining biological diversity (Vellend 2016). Both niche and neutral processes provide a solid theoretical background to model species assembly into local assemblages (Gravel et al. 2006) distributed in a metacommunity context.

Consider a metacommunity *M* defined by sets of local assemblages (*k*) with some degree of connection by dispersal. Let *N_*k_* be the number of individuals inhabiting each *k* assemblage, and *N_*M_* be the total number of individuals in the metacommunity, such that 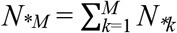. In the absence of a major disturbance, *N_*k_* is expected to increase until *k* achieves carrying capacity (*J_k_*), quantified as the maximum number of individuals allowed to inhabit *k*, given the minimum amount of space and resources per capita demanded by the organisms (Del Monte-Luna et al. 2004). Thus, under *Jk*, we expect *dN_*k_*(*t*) = 0, and species assembly likely starts to follow a zero-sum dynamic (Hubbell 2001). This means that an individual currently inhabiting *k* must necessarily die before a new one can be recruited, either by birth or immigration from other assemblage.

Given there is vacant space in *k* to be occupied by new recruits, what processes will determine their identity? As we seen before, niche processes may be the answer to this question, at least partially. For a given species *i* belonging to the species pool *q*, the niche position *y,* can be defined as the environmental condition that maximizes the probability of *N_ik_* > 0 such that *dN_ik_*(*t*) ≥ 0. Further, the niche breadth *s_i_* can be defined as the amount of deviation from *y_i_*, that allows the conditions *N_ik_* > 0 and *dN_ik_*(*t*) ≥ 0 to be satisfied (Sexton et al. 2017). From a niche perspective, and considering only the influence of environmental filtering, the probability (*λ_ik_*) of species *i* to recruit a new individual in *k* can be modeled as a deterministic Gaussian function based on their niche position and breadth, as follows:

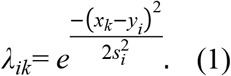

Accordingly, *x_k_* is the environmental position of *k* across an environmental gradient, *y_i_,* is the niche position of *i* (measured in the same scale of *x_k_*), and *s_i_*, is the niche breadth of *i*. This model has been originally proposed by Minchin (1987), and has been adopted in several studies to simulate species assemblages (Dray and Legendre 2008; Duarte et al. 2018; Nakamura et al. 2020; Peres-Neto et al. 2017; Peres-Neto et al. 2012; Sokol et al. 2017; Sokol et al. 2015).

We may now compute a recruitment probability (*R*_Niche_*ik*__) that implicitly incorporates the effect of biotic interactions (Gravel et al. 2006):

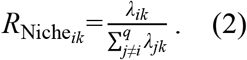

Note that *λ_ik_* is standardized by the probabilities of all other *j* species, such that *j* ∈ *q*, which are also candidates to recruit new individuals, competing directly with *i.* Since *R*_Niche_*ik*__ is only defined by niche position and breadth, *q* can contain not only the species already resident in *k*, but also potential immigrant species occurring at any assemblage *l,* such that *l* ∈ *M*. The only condition to be a candidate recruiter species to *k* is to show *R*_Niche_*ik*__ > 0. This condition agrees with the species sorting model of metacommunity theory (Leibold et al. 2004).

Gravel et al. (2006) improved this niche-based model of species assembly by incorporating parameters from Hubbell’s neutral model (Hubbell 2001), which provides spatial context to species assembly in local assemblages. First, let us consider dispersal limitation as a mechanism affecting the success of any species *i* belonging to the regional species pool *q*, and occurring in any *l* assemblage, to disperse new recruits to assemblage *k*. The probability *δ_lk_* of any *i* inhabiting *l* to contribute to the immigrant pool of *k* may be modeled as a Gaussian function of the shortest linear distance (*r,* rescaled between 0 and 1, see Sokol et al. 2017) between *k* and *l*, and the slope *w* of the dispersal kernel (the probability density function of the dispersal success from a source to a sink assemblage, see Nathan and Muller-Landau 2000) for the species in *l* (Gravel et al. 2006; Sokol et al. 2017; Sokol et al. 2015), as follows:

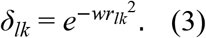

For a slope *w* = 0, *δ_lk_* = 1, independently of the distance between *l* and *k*, which means that the species in assemblage *l* face no barrier to disperse to *k* (the assemblages are panmictic). On the other hand, for slope *w* = 10, *δ_lk_* = 0.9 for sites showing *r* = 0.1, but decreases fast to *δ_lk_* = 0.08 if the distance between the assemblages increases to *r* = 0.5. Over the next sections we introduce a simulation-based framework to estimate *w* for empirical datasets, which enables quantifying dispersal limitation underlying diversity patterns (see *Estimating niche and neutral parameters underlying species diversity gradients using Approximate Bayesian Computation*).

The probability of any species *i* to successfully disperse new recruits from any assemblage *l* (*R*_Dispersala_*iM*__), is a function of the abundance of *i* across *M*, or 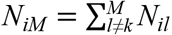, and *δ_lk_*, and is contingent on the probability of dispersal for all other species *q* in *M* (Gravel et al. 2006), as follows:

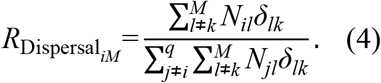

It is important to observe that even for *δ_lk_* = 1, immigrant species arriving to site *k* may not succeed as new colonizers, depending on how resistant *k* is to immigration. Assemblages where resident species are strong recruiters may prevent the entrance of new colonizers (e.g. Duarte et al. 2006), which reduces the chance of new species become part of the assemblage, even in the absence of dispersal limitation. For a site *k* under zero-sum dynamic, let *d*_k_ be the number of instantaneous deaths in *k*. Site *k* is now open to the recruitment of *J_k_* – *d_k_* new individuals, which may be generated either by resident parents inhabiting *k* (*b_k_*), or by individuals dispersing from any site *l* ∈*M* (*b_l_*), such that *J_k_* = (*b_k_* – *d_k_*) + *b_l_*. The immigration coefficient *m_k_* (Hubbell 2001) can then be computed as

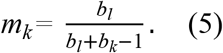

Now we can compute the recruitment probability of a species *i* in the assemblage *k* given the neutral parameters *R*_Dispersa_*iM*__, *N_ik_, N_iM_*, and *m_k_* (Sokol et al. 2015), as follows:

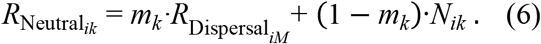

Finally, we compute the recruitment probability of any species *i* ∈ *q* in any assemblage *k* ∈ *M* that considers both niche and neutral processes, as

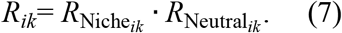

The definition of *R_ik_* provides an integrative solution to model species assembly patterns in local assemblages that allow estimating the strength of niche and neutral process determining the dynamics of biological diversity across space. Yet, the evaluation of the imprint left by niche evolution on species assembly, and ultimately on biological diversity, is still lacking. Over the next section we present a solution to fill this gap.

### Niche evolution and species assembly across environmental gradients

The Ornstein-Uhlenbeck (OU) model of trait evolution (Hansen 1997) allows parameterizing the strength of the adaptation rate α of species niches evolving towards an optimum niche condition θ, as follows:

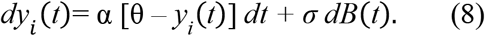

For any θ and given the niche position *y_i_*(*t*) of species *i*, adaptation rate *α* is a force that deterministically decreases the fitness of *y_i_*(*t*) as θ – *y_i_*(*t*) ≠ 0. As α → 0, the strength of adaptation rate decreases, and trait divergence among species is increasingly modeled by the stochastic component of the OU model (*dB*[*t*]), which is a stochastic white-noise process showing Gaussian distribution (*N*[0,*dt*)], Brownian motion), and standard deviation σ, which indicates the amplitude of niche position variation among species (Butler and King 2004; Hansen et al. 2008). Therefore, under low α values, species niche position evolves under strong phylogenetic inertia, generating high phylogenetic signal in species niche positions (Blomberg and Garland 2002). On the other hand, as α → ∞, species niches tend to evolve faster from the ancestral state (*yi*_anc_.) towards θ, and the imprint of phylogenetic inertia on species niches tends to be erased. The rate of niche evolution towards θ can be estimated as a phylogenetic half-life (Hansen 1997; Hansen et al. 2008), which indicates the amount of time units needed to move half the distance between the ancestral state and the optimum condition, and is calculated as

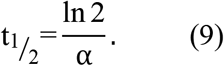

Note that *t*_1/2_ is expressed in time units, which means that the effect α exerts on niche evolution depends on the timespan between *yi*_anc_. and θ. If the timespan is long enough (say, 200 million years), even a small α value (say, α = 0.01) may generate a moderate movement towards θ (*t*_1/2_ = 69.3). Nonetheless, if the evolutionary timespan is shorter than ≡ 70 million years, the phylogenetic half-life corresponding to α = 0.01 will be too slow to produce significant niche evolution towards the niche optimum, and therefore niche evolution may show phylogenetic inertia not quite distinguishable from Brownian motion (α = 0, *t*_1/2_ = + ∞). Thus, the expected value of a niche *y_i_*, evolving under OU model during a time *t*, results from a deterministic process modulated by the adaptation rate α that forces *y_i_*(*t*) to converge towards the optimum niche condition θ, and a stochastic, Brownian process, as follows:

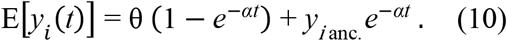

The critical step is to translate the OU parameters α and θ into quantities that can be estimated from assemblages described by species abundances or occurrences, an environmental variable associated with species assembly, and a phylogenetic tree. In a metacommunity *M* composed by *k* local assemblages defined by *q* species belonging to a monophyletic lineage, those local assemblages with the highest levels of biological diversity can be considered as offering the best environmental condition *x* to most species in *q* (Enquist et al. 2015; Violle et al. 2007), and therefore it is likely to be an appropriate estimator of θ. We might then take *x* as an empirical estimate of θ, or *x*_θ_.

Niche filtering is expected to be stronger in assemblages with environmental condition deviating from *x*_θ_, leading to lower species diversity. Differently from *x*_θ_, which can be estimated from data, the parameter α cannot be directly assessed, but its posterior distribution can be estimated by means of rejection-based Approximate Bayesian Computation (Beaumont 2010; Csilléry et al. 2010). We explain this in detail in the next section.

For now, it is important to understand what the parameter α can inform about the role of niche evolution on species assembly patterns. If α → 0, we may conclude that niche evolution was mainly a stochastic Brownian process, which means that the adaptation rate towards *x*_θ_ was very weak along niche evolution, insufficient to break the phylogenetic signal of the ancestral niche of species. Under such evolutionary scenario, niche divergence between species will be a function of the time since their evolutionary divergence, leading to strong phylogenetic covariation in species niches. Furthermore, the total variance in the niche position will be high, which means that niche positions of species will be more loosely distributed across the upper and lower limits of the environmental condition *x*. Niche-based assembly processes in local communities will tend to be phylogenetically structured, while differences in diversity values among assemblages will be not remarkable. As α → ∞, adaptation rate of species niches towards *x*_θ_ becomes faster along niche evolution, so that any track of phylogenetic signal in the niche vanishes along evolution (Cooper et al. 2016). Niche divergence between species will be constrained by the optimum niche condition, with values approaching *x*_θ_ as α increases, decreasing the total variance of the niche position among species. In local assemblages showing environmental conditions close *x*_θ_, steep diversity peaks are expected to occur.

### Estimating niche and neutral parameters underlying species diversity gradients using Approximate Bayesian Computation

Consider that assemblages in *M* are distributed across an environmental gradient defined by a single environmental condition, or multiple conditions summarized in a single factor or principal component *x*. We may estimate *x*_θ_ as the environmental condition interval that maximizes species diversity in *M*. In turn, whenever the niche position *y_i_* of a given species in *M* approaches *x*_θ_, the probability of occurrence of that species in the assemblage increases, conditioned to the niche positions of all other species in the assemblage. The adaptation rate of *y* along the evolutionary history (i.e. OU’s α) is a niche-assembly parameter that can be estimated given the species diversity variation in *M*, *x*, the spatial distribution of local assemblages in *M*, and the phylogenetic relatedness among species distributed across *M*. By doing so, we can reveal how strong is phylogenetic signal in niche position of species. On the other hand, species diversity in a local assemblage may be limited by the dispersal capacity of species. As we have seen before, this depends on the spatial distance between sites (*r*) and the slope *w* of the dispersal kernel, in complement to niche characteristics of species (Gravel et al. 2006). Therefore, estimating *w* allows quantifying the influence of dispersal limitation on the distribution of species diversity across *M*.

To estimate niche (OU’s α) and neutral (slope *w*) parameters associated with the distribution of species diversity, we developed an Approximate Bayesian Computation (ABC) algorithm (Beaumont et al. 2002; Csilléry et al. 2010) that works as schematically represented in figure 1.

**Figure 1:**
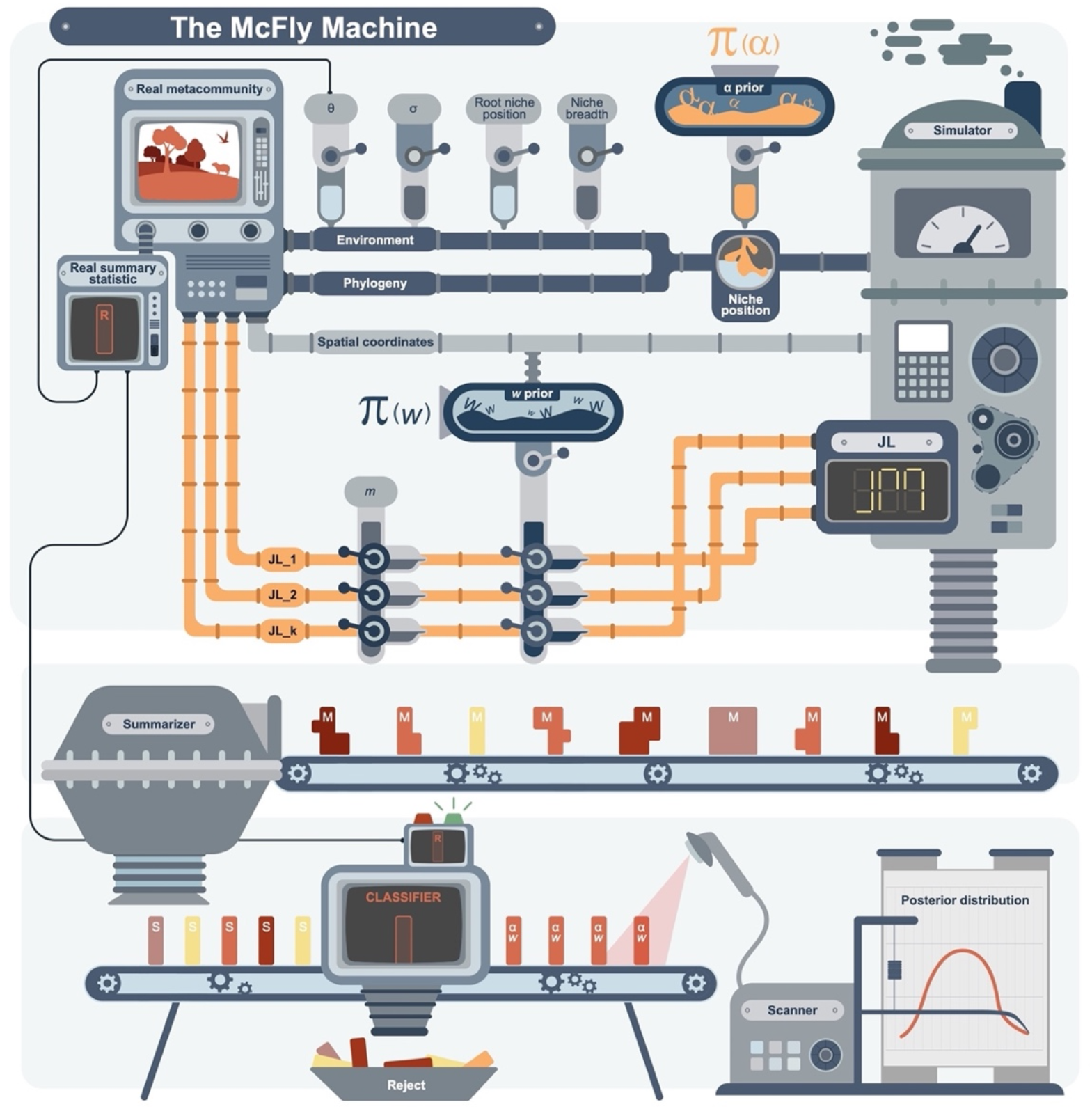
Schematic representation of *mcfly,* an Approximate Bayesian Computation framework developed to estimate the posterior distribution of adaptation rate (OU’s α) and dispersal kernel slope (*w*) parameters driving species diversity gradients along an environmental gradient. **1**. The *real metacommunity* provides the empirical measures of species diversity, the *real summary statistic (R).* **2**. *Environment* is a dimension of species niches that provides the optimum niche (θ), variance (σ) and root niche position parameters necessary to simulate species niche based on the *phylogeny,* according to Ornstein-Uhlenbeck evolutionary model. We also define niche breadth in the simulation based on *environment.* **3**. A prior distribution of OU’s α is used to simulate species niche positions based on *environment* parameters and *phylogeny.* **4**. *Spatial coordinates* of *k* sites are used to define the prior distribution of slope *w*; together with immigration rate (*m*), number of individuals at *k* sites (*J_L_k_*) and the total number of individuals in *M*(*J_M_*) provide the neutral parameters of simulations. **5**. Metacommunities are simulated based on niche and neutral parameters (*Simulator*). **6**. Each metacommunity simulation provides species diversity measures (*Summarizer*) taken as the simulated summary statistics (*S*). **7**. Simulated diversities (*S*) are classified based on a pre-defined minimum correlation threshold (tolerance) to the observed diversity (*R*). Every *S* object not discarded (*Reject*) is then *scanned* to record its OU’s α and slope *w* to provide a posterior distribution of adaptation rate (α) and dispersal limitation (*w*), respectively.

We implemented the ABC framework in the R package *mcfly* (freely available at (https://github.com/GabrielNakamura/mcfly), which has ‘mcfly’ as the core function. An additional function that helps users to define the appropriate tolerance threshold (‘define_tolerance’) is also available in the package, as well as a working example to illustrate how the user can set all the parameters needed to run ‘Mcfly’ function. The example can be easily accessed in R by typing *vignette(“mcfly_vignette”).*

### The ‘simulator engine’

The simulation framework implemented in *mcfly* package derives from the individual-based simulation protocol developed by Sokol et al. (2015, 2017) and available in the R package *MCSim* (available at https://github.com/sokole/MCSim). Accordingly, sets of assemblages are simulated based on lottery colonization and recruitment dynamics, following niche and neutral mechanisms (Gravel et al. 2006), as explained in the section *Parameterizing niche and neutral processes determining species assembly*.

#### Metacommunity dimensions and species pools

Simulated metacommunities *M_Sim_* show the same number of sites of the observed metacommunity *M*. If *M* contains information on species abundances, the number of individuals in each simulated assemblage equals the number shown in the respective observed local assemblage. Because the original simulation protocol of *MCSim* is individual-based, when only species incidence (presence/absence) data is available the simulation performed in *mcfly* addresses 100 individuals to each species present in each local assemblage prior to the simulation of *M_Sim_*. In the end, *M_Sim_* is transformed to an incidence matrix.

Noteworthy, the phylogenetic tree used to build *M_Sim_* should be as complete as possible to increase the quality of niche simulations, which means it should not be pruned to keep only the species present in *M.* In all simulations, the species pool of *M_Sim_* will be the same defining *M,* independently of the size of the species pool included in the phylogeny. According to Sokol et al. (2015), recruitment patterns in *M_Sim_* stabilize after 30 simulation time steps. By default, we set simulations to stop after 50 timesteps.

#### Niche prior and other parameters

At each simulation run, the niche position of each species is simulated for all species included in the phylogenetic tree using the function ‘rTraitCont’ of the package *ape* (Paradis and Schliep 2019) as a continuous trait *y* following OU model. The adaptation rate parameter α of niche simulation is randomly sampled from a prior distribution, with predefined size and distribution mode. The minimum value of the α parameter is set to correspond to the half-life (t_1/2_, see eq. 9) equal to the total timespan of the phylogeny. For a phylogeny with timespan = 30 Myr, the minimum α value will be 0.023, since

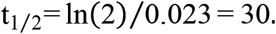

The maximum α parameter value corresponds to t_1/2_= 1/30 · timespan. Thus, for that same timespan = 30 Myr, the maximum α value = 0.693, since

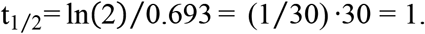

In the example above, the range of α parameters between the minimum and maximum limits expresses a broad variation in phylogenetic signal. Indeed, we observe an asymptotic decay in phylogenetic signal, estimated by Blomberg’s K (Blomberg and Garland 2002), with increasing OU’s α values (fig. 2). Considering the α parameter interval ranging between 0.023 and 0.693, K statistics is shown to vary between 0.983 and 0.137, respectively. Further, the lower 95% credibility limit for OU’s α values estimated from vectors randomly drawn from Gaussian distribution (white noise) reveals that niche positions showing α values higher 0.628 cannot be distinguished from white noise variables and show very low phylogenetic signal ≤ 0.165. Therefore, we conclude that the range of prior α values defined in the simulation algorithm is appropriate. Note that the simulation does not rescale the branch lengths of phylogenetic tree. Therefore, the prior distribution of OU’s α values will differ from the above whenever the timespan differs from 30 Myr, and therefore the interpretation of results must consider the timespan of the phylogenetic tree under analysis. By default, *mcfly* generates a uniform distribution of the α prior, but the user may opt for generating a prior α with half-life values uniformly distributed, instead. In the next sections we provide a sensitivity analysis to evaluate the implications of the α prior distribution for the posterior distribution of OU’s α parameter.

**Figure 2:**
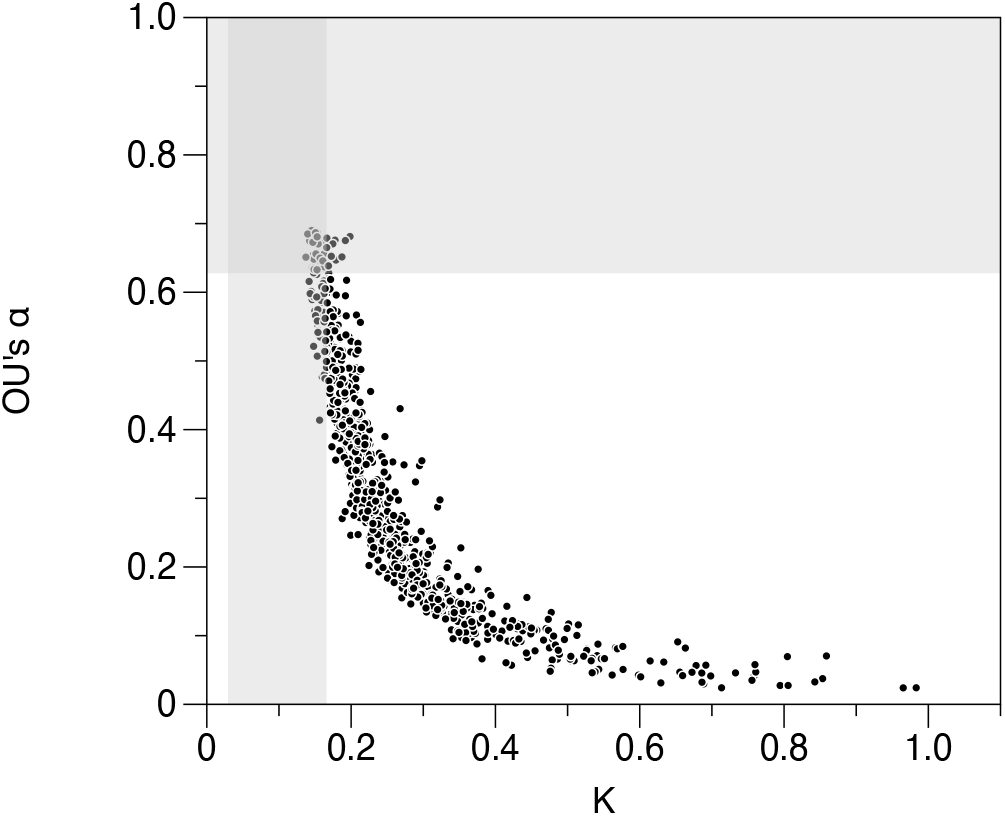
Relationship between Blomberg’s K statistic and the adaptation rate parameter of Ornstein-Uhlenbeck model (OU’s α) for simulated niche positions computed for 1,000 sets of 50 species each (black circles). Gray areas indicate lower and upper credibility intervals (95%) for Blomberg’s K and OU’s α values estimated from white noise vectors (*N*[0.1]).

Another critical niche parameter to be defined is the optimum niche position (*x*_θ_). In our framework, *x*_θ_ is empirically defined as the values of the environmental condition *e* showing the highest species diversities in *M*. In *mcfly*, all *e_k_* values corresponding to ≥ the 5% highest species diversity values will constitute a vector of *x*_θ_ values. At each metacommunity simulation process, a single *x*_θ_ value is randomly drawn from that vector and taken as the optimum niche. Thus, it is important to note that the more unimodal is diversity distribution across *e*, the more homogeneous will *x*_θ_ among different *M_Sim_*, which is likely to decrease the credibility intervals of the posterior distribution of OU’s α and slope *w* parameters. Moreover, the variance of the niche position (σ) was empirically defined as 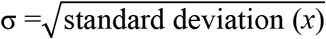, and root node value = 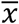.

#### Neutral prior and other parameters

The spatial coordinates of each assemblage in *M* are used to compute pairwise distances *r* between local assemblages in *M*, which are rescaled to vary between 0 and 1. As we have seen in equation 3, *r* is a component defining the probability of a species to disperse from a site to another, whose effect on dispersal limitation is modulated by the slope *w,* the neutral assembly parameter of interest in our framework. It is important to note that the range of variation of *w*, which expresses the strength of dispersal limitation, depends on the shape of the spatial distribution of sites. When sites are evenly distributed across the space, as in a cell grid, *w* slopes must vary over broader scales to capture dispersal limitation over shorter distances, and vice-versa. Thus, to build a prior distribution of slope *w*, we first defined the maximum distance *r* between sites enough to connect all sites in *M* by means of a minimum spanning tree (Legendre and Legendre 2012). This threshold distance provides information on how far two sites should be to each other to dispersal limitation to become relevant and is widely used in spatial analysis (Borcard and Legendre 2002). Then, we simulated a uniform distribution of dispersal probabilities between sites (δ_lk_, see eq. 3), varying between 0 and 1, and with a predefined size. Based on equation 3, we then computed the slope *w* corresponding to every simulated δ_lk_ for a hypothetical pair of sites located at distance *r* to each other. By doing so, we build the prior distribution of slopes *w*, which is not uniform itself, but decays logarithmically as *w* increases.

As we have seen in eq. 5, the probability of immigration in each assemblage (*m*) has also to be defined prior to the simulation, which is not trivial in most cases. The user may opt for indicating an arbitrary value; the default value in *mcfly* is 0.5, which means that the simulation allows a half of the individuals to be replaced at each timestep of the simulation process. Alternatively, the user may inform empirically estimated *m* for each site, whenever these data are available.

#### The summary statistic

Our simulation framework summarizes species diversity in the local assemblages of *M* and *Msim* using Rényi entropy (Rényi 1961), which is a flexible measure that generalizes widely used diversity measures, such as Shannon and Simpson diversity indices (Anand and Orlóci 1996). Rényi entropy (H_*order*_) is computed as follows:

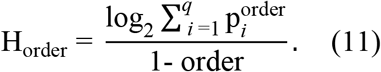

Accordingly, *q* is the number of species present in the community, *p*_i_ is the proportional abundance of each species *i* in the community, and order is a scalar that modifies the sensitivity of entropy to variation in species abundances (evenness). In this study, we analyzed three entropy orders: 1, 2, and 12. For entropy order = 1, Rényi entropy is equivalent to Shannon diversity index, while entropy order = 2 assumes values equivalent to Simpson index (Anand and Orlóci 1996). Higher entropy orders increase the importance of evenness in the diversity index, which tends to stabilize for entropy order > 12 (Anand and Orlóci 1996). For datasets characterizing sites by species incidences (presence/absence data), Rényi entropy is equivalent to species richness (or entropy order = 0), independently of the entropy order used.

#### The posterior distribution of adaptive rate (α) and dispersal limitation (w)

Since species diversity is computed for *M* and all *M_Sim_* datasets, the correlation between observed and simulated diversities is a way to evaluate how similar are observed and simulated metacommunities. This step involves defining a threshold correlation value below which metacommunities are rejected. In ABC analysis, a tolerance value is usually defined, and expresses the maximum distance tolerated between observed and simulated summary statistics (Beaumont 2010). In *mcfly*, the tolerance expresses the complement of the correlation (1 – correlation) and must be informed prior to the simulations. The remaining *M_Sim_* datasets surpassing the tolerance threshold provide their corresponding OU’s α and slope *w* parameters to a posterior distribution of niche and neutral parameters underlying diversity patterns in *M*, respectively. A highest posterior distribution (HPD) provides the credibility intervals of those parameters (Turkkan and Pham-Gia 1993).

### Evaluating *mcfly* performance using simulated metacommunities

To evaluate the performance of *mcfly* we simulated 2,000 metacommunities *M* using the function ‘metasim’ of the package *MCSim* (Sokol et al. 2017; Sokol et al. 2015), each of them showing 50 local assemblages, with known niche and neutral parameters, which allowed to evaluate the efficiency of the proposed simulation approach to estimate them.

Prior to metacommunity simulation, phylogenetic trees containing 50 species were simulated using the function ‘sim.bdtree’ of package *geiger* (Harmon et al. 2008), assuming homogeneous birth-death processes across all lineages. Speciation followed purely stochastic birth (speciation rate = 0.1, extinction rate = 0). The number of lineages increased exponentially with time. Evolutionary timespan was fixed as 30 Myr. These phylogenies defined the species pools used to perform all further simulations. All 50 species in each phylogeny were allowed to occur in at least one out of 50 communities representing a metacommunity *M*.

For each phylogeny, a single OU’s α value defining the real adaptive rate of species niche positions was randomly drawn from a uniform distribution ranging between minimum (= 0.023) and maximum (= 0.693) α values. Further, an environmental variable *x* was simulated from a uniform distribution (U[0,100]), and its mean value 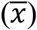 was defined as the optimum niche position (θ). OU’s α, 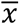 and other niche parameters described in the section *Niche prior and other parameters* were then used to simulate the niche positions *yi* of species ranging between 0 and 100 based on OU method of the ‘rTraitCont’ function of package *ape* (Paradis and Schliep 2019). For metacommunity simulation, species niche breadth (*si*, eq. 1) was fixed as 10, which confers reasonable niche breadth to species (Duarte et al. 2018; Peres-Neto et al. 2012).

Spatial coordinates of assemblages in *M* were simulated from a uniform distribution (U[0,1]). The number of individuals per local assemblage and the total number of individuals in *M* were defined as, respectively, *J_k_* =1000 individuals, *J_M_* = 1000 x 50 sites = 50,000 individuals. The probability of immigration (*m,* see eq. 5) was defined as 0.5. The frequencies of species across *M* were extracted from a multinormal distribution ranging between 0.1 and 0.4 that was further normalized based on *J_M_*. Species colonization or recruitment across *M* stopped after 50 simulation time steps. Noteworthy, in some simulations, a low number of species was capable of colonizing at least one local assemblage in *M*, and therefore gamma diversity in *M* varied among the simulations. We analyzed only *M* datasets containing, at least, 20 species distributed across local assemblages. We divided the 2,000 *M* datasets into five subgroups of 400 metacommunities, each of them simulated under different dispersal limitation strengths (*w* = 0, 1, 5, 15 or 100, see *Neutral prior and other parameters*). Thus, the real *w* slope of each simulated *M* was known. After metacommunity simulation, each *M* dataset was converted into an incidence matrix describing species occurrences in each local assemblage.

After defining *M* datasets, we submitted each metacommunity to ABC analysis using *mcfly* to evaluate the suitability of the framework to estimate the posterior distribution of the adaptive rate of niche positions (OU’s α) and dispersal limitation strength (slope *w*). Prior to ABC analysis, the 2,000 metacommunities *M* were split into two groups containing 1,000 metacommunities each. In the first group, the prior distribution of OU’s α used in the ABC algorithm was defined as a uniform distribution of α values (‘uniform’ α prior). In the second group, α prior distribution was defined based on a uniform distribution of α half-lives (‘half-life’ α prior), and the corresponding α values were then taken as OU’s α prior (see *Niche prior and other parameters*). The maximum sample size of α and *w* priors used to analyze each metacommunity *M* was defined as 14,400 simulations of *M_Sim_* datasets, while the posterior sample size was defined as 240. Since we had no previous information about the appropriate tolerance value (see previous section), prior to ABC we computed correlations between the summary statistics computed for each *M* dataset and for 480 *M_Sim_* datasets to obtain a correlation distribution. The complement of the 95^th^ percentile of the respective correlation distribution was then taken as the tolerance value in ABC performed for each metacommunity *M*.

At each simulation step, *mcfly* simulated *M_Sim_* under predefined adaptation rate (OU’s α) and dispersal limitation (slope *w*), computed the summary statistics (Rényi entropy of order 1), and correlated it to the real summary statistics computed from *M*. Whenever the complement of correlation was ≤ the tolerance limit, α and *w* parameters were recorded in the posterior distribution of α and *w* parameters. After the algorithm reached either the posterior sample size or the maximum prior size (in cases where the posterior sample size was not reached), the median values of each parameter in the posterior distribution were recorded.

### Statistical performance of ABC based on simulated datasets

We evaluated the statistical performance of the ABC approach using structural equation modelling based on *d*-separation tests (Shipley 2000), which estimates the validity of causal models using Fisher’s C as test statistic that follows a chi-square distribution with 2*k* degrees of freedom, where *k* is number of independence relationships necessary to evaluate the model given the number of variables and the number of causal relationships proposed in the model (Shipley 2000). Accordingly, the probability of the null hypothesis of the fitted model to correspond to expected causal relationships indicates the overall validity of the proposed model. Path coefficients (*p*) linking variables in the causal model were estimated using ordinary least squares (OLS). The analysis was performed using the ‘psem’ function of the package *piecewiseSEM* (Lefcheck 2016).

We built the causal models considering first the effects of both real OU’s α and *w* on the number of species in *M* (which varied across simulations, as some species were lost during the assembly), and on the median of the posterior distribution of both OU’s α and *w*. Furthermore, we evaluated the influence of the prior OU’s α mode (‘uniform’ or ‘half-life’) on the median of the posterior distribution of OU’s α. Finally, the influence of the number of species in *M* on both was the median of the posterior distribution of both OU’s α and *w.* In a second set of structural models, we replaced real and posterior OU’s α by the respective half-life and tested similar causality relationships. Prior to causal testing, half-life and *w* slopes were log transformed.

We also evaluated causal relationships within each prior OU’s α mode. These models were tested because we found that the effect of prior OU’s α mode on the median of the posterior distribution of OU’s α was stronger than the effect of the real OU’s α (the same occurred in the half-life model). Evaluating causal relationships within each prior OU’s α mode enabled us to perform a sensitivity analysis of the ABC approach to the prior OU’s α.

In all causal models, we considered that the residuals of the median of the posterior distribution of OU’s α and *w* slope were correlated to each other.

### A study case: phyllostomid bats

We analyzed the influence of niche evolution and dispersal limitation on the distribution of species diversity of the Phyllostomidae, a species-rich and ecological diverse family of bats. Phyllostomid bats are predominantly distributed in the Neotropics, from California to northern Argentina and Chile (Fleming et al. 2020). The family has 222 known species (Simmons and Cirranello 2020), displaying a large array of dietary and morphological specializations to feed on fruits, nectar, and other parts of plants, as well as on insects, vertebrates, and blood (Baker et al. 2012; Fleming et al. 2020; Monteiro and Nogueira 2011; Rossoni et al. 2019). The remarkable diversity of their ecological features mirrors a strong latitudinal and environmental gradient of species richness (Pereira and Palmeirim 2013; Villalobos and Arita 2010), which makes this family particularly appropriate for our purpose. We gathered geographic range maps for phyllostomid species from IUCN database (IUCN 2020), and defined species assemblages based on species incidences in a grid composed by 110 × 110 km cells distributed across the Neotropical region strictu sensu (see Maestri and Duarte 2020; Morrone 2014). Phylogenetic relationships among species were taken from the consensus phylogenetic hypothesis proposed by Shi and Rabosky (2015), the most comprehensive bat phylogeny available to date. The complete Chiroptera phylogeny was dated using 24 fossil calibration points distributed along the whole timespan of the tree. Pruning geographic data given available genetic information resulted in a dataset of 136 species (*ca.* 62% of total phyllostomid richness) distributed across 1,338 assemblages in the Neotropical region. The total timespan of the pruned phylogeny was = 34 Myr.

We computed entropy of order 1 for sites described by phyllostomid species and analyzed its association with each of four climatic variables (mean annual temperature, temperature seasonality, mean annual precipitation and precipitation seasonality) and two topographical variables (mean altitude and altitudinal range variation). Climatic variables were taken from the WorldClim2 database (Fick and Hijmans 2017), and topographic variables were taken from NASA (http://neo.sci.gsfc.nasa.gov/). Phyllostomid entropy was strongly negatively related to temperature seasonality (*r* = - 0.91), corroborating results obtained elsewhere (Stevens 2011), and was taken as the niche condition (*x*) in further analysis.

Then, based on the species by sites matrix (assemblages), the spatial coordinates of the sites, the niche condition (precipitation seasonality), and phylogenetic relationships among species, we were able to estimate the posterior distribution of the adaptive rate of species niche positions (OU’s α) and dispersal limitation strength (slope *w*) underlying the distribution of species diversity across phyllostomid assemblages using ABC analysis implemented in *mcfly*. To simulate niche positions, we defined 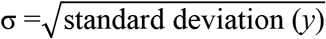, root node value = 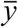. The probability *m* of new colonizers to arrive at simulated assemblages = 0.5. Niche breadth (*s_i_*) was set to 10. Species colonization or recruitment in each set of simulated communities stopped after 50 simulation time steps. We set the maximum sample size of OU’s α (uniform mode) and *w* priors to 36,000, and the size of the posterior sample to 480. The tolerance in ABC analysis (1 – *r*) was set to 0.1.

## Results

### Simulated datasets

The results of global causal model analyzing factors influencing the posterior distribution of OU’s α and *w* slope are shown in figure 3*A*. The overall model fit (Fisher’s C = 2.67, *P*_df=4_ = 0.62) indicated that the model was causally valid. The median of the posterior distribution of OU’s α was mostly explained by the OU’s α prior mode (*p* = 0.69), and to a lesser extent by the real OU’s α (*p* = 36) and the real w slope (*p* = 0.2). The number of species in the metacommunity did not affect the median of the OU’s α posterior distribution (*p* < 0.01). On the other hand, the median of the posterior distribution of *w* was mostly explained by the real *w* slope (*p* = 0.85). Replacing the median of the posterior distribution of OU’s α by half-life (fig. 3*B*) did not change the overall architecture of causal relationships. The overall model remained valid (Fisher’s C = 1.18, *P*_df=4_ = 0.88); noteworthy, the strength of the path coefficient linking the real half-life to the median of the posterior distribution of OU’s α increased (*p* = 0.42), while the latter was still mostly explained by the OU’s α prior mode (*p* = - 0.74). All other causal relationships behaved similarly to the observed in the previous model (fig. 3*A*).

**Figure 3:**
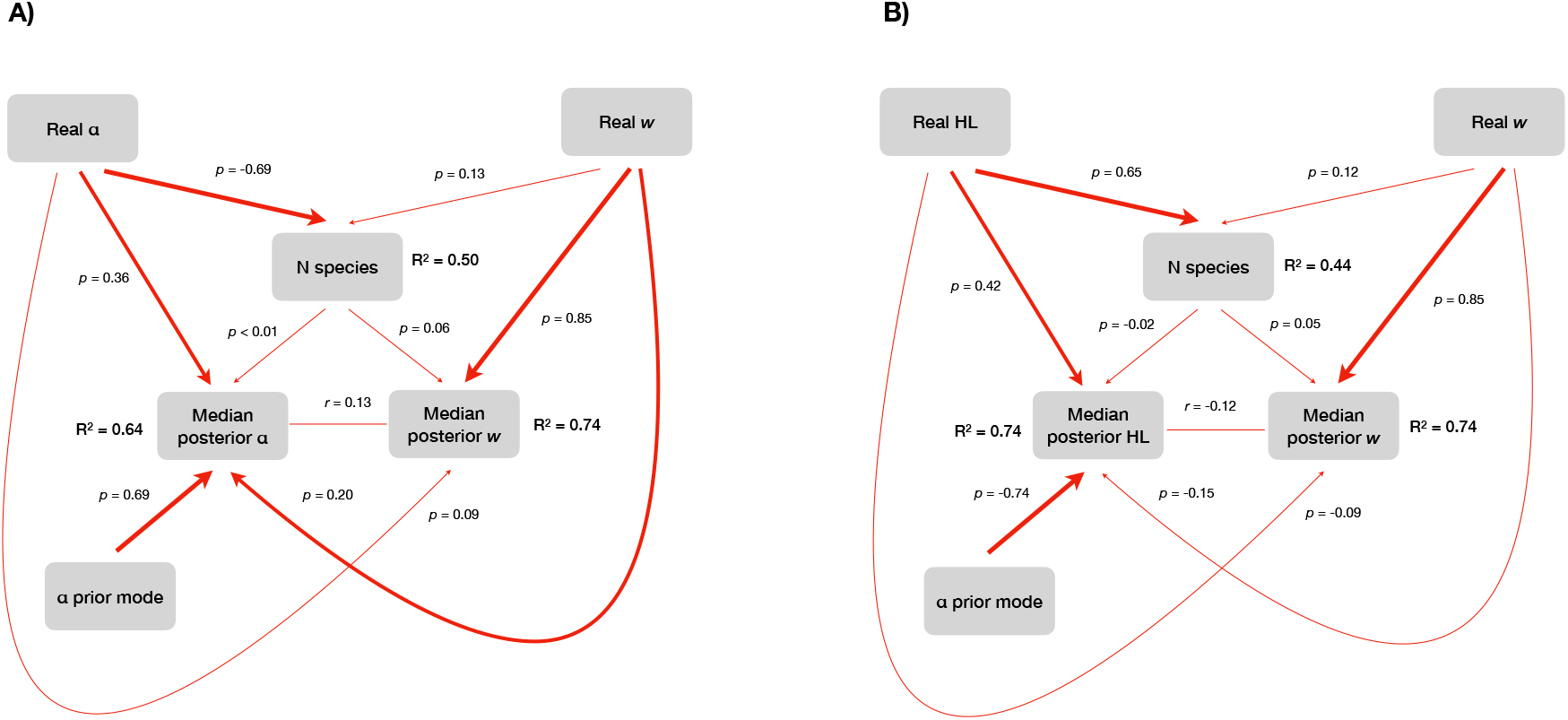
Causal models relating real OU’s α or half-life, real *w* slope, OU’s α prior mode and number of species in 2,000 simulated metacommunities on the median of the posterior distribution of OU’s α of half-life and *w* slope. Causal relationships were estimated using ordinary least squares. Half-life and *w* slopes were log-transformed prior to analysis. Validity of overall models was tested using d-separation tests (Shipley 2000). *A*) OU’s α model (Fisher’s C = 2.67, *P*_df=4_ = 0.62). *B*) Half-life model (Fisher’s C = 1,18. *Pdf=4* = 0.88). Arrow width indicates the strength of the respective causal relationship.

Bivariate relationships among real OU’s α, half-life and *w* slope and the corresponding median values of the posterior distribution of OU’s α, half-life and *w* slope are shown in figure 4. In figures 4*A, C* and *E*, we observe relationships obtained when the prior distribution of OU’s α was uniform, whereas figures 4*B, D* and *F* show the relationships obtained using the half-life derived prior distribution of OU’s α. Exploring these bivariate relationships suggests that using the uniform prior distribution of OU’s α produce stronger associations between real OU’s α or half-life and the median posterior distribution of OU’s α (fig. 4*A*) or half-life (fig. 4*C*), respectively, than the corresponding relationships based on half-life derived OU’s α prior (figs. 4*B, D*). But even using the uniform OU’s α prior, the dispersion of the median posterior distribution of both OU’s α and half-life tended to increase towards higher real OU’s α, which should possibly lead to lower reliability of the posterior distribution of the α parameter under higher real OU’s α. Such tendency was minimized when the association between log-transformed real half-life and median of the posterior distribution of half-life was considered, as we also found a linear relationship between those variables when the uniform prior distribution of OU’s α was used (fig. 4*C*). Further, the association between real *w* and the median posterior distribution of *w* slope was not affected by OU’s α prior mode, as expected (figs. 4*E*, *F*).

**Figure 4:**
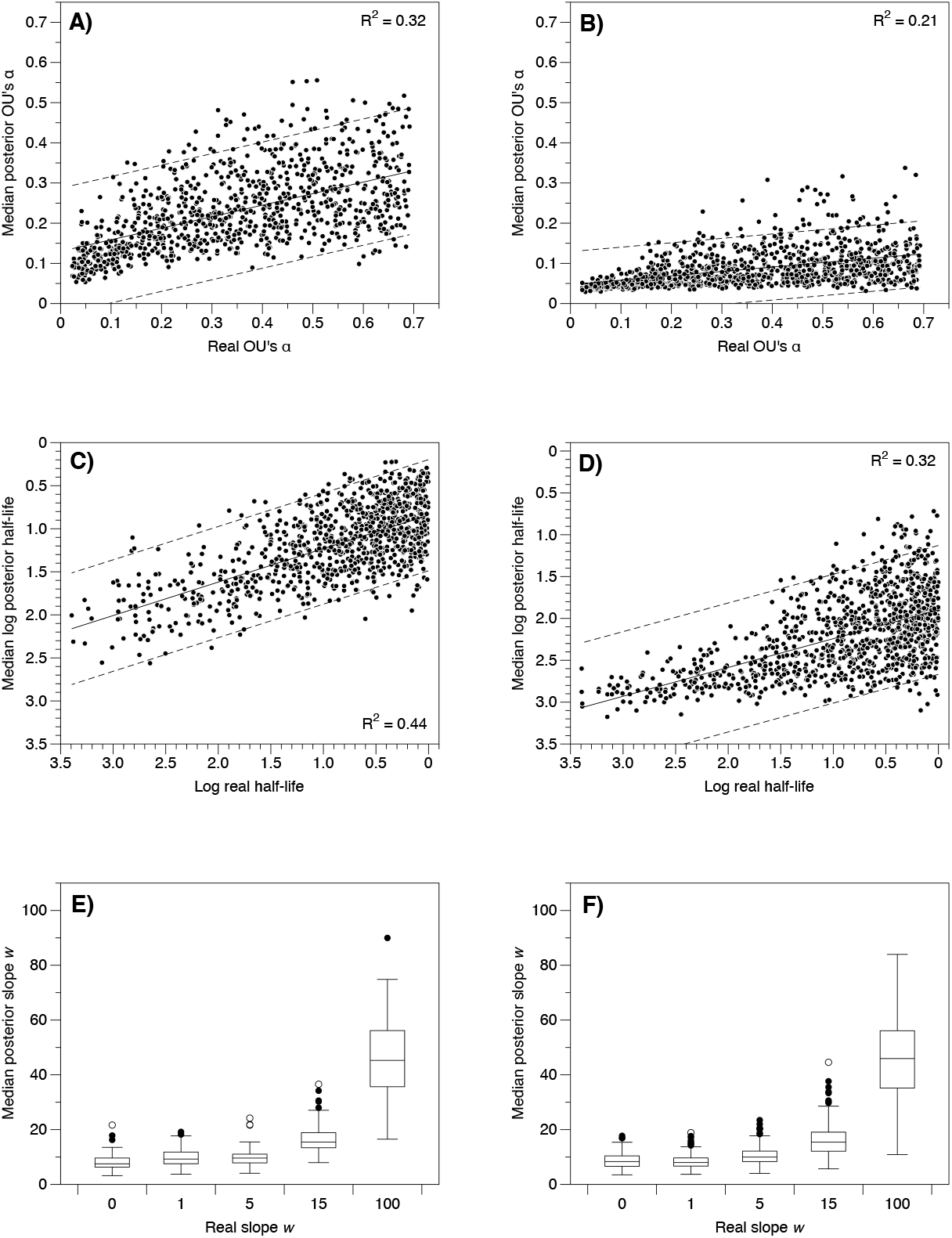
Bivariate relationships between real niche (OU’s α or half-life) or dispersal (*w* slope) parameters of simulated metacommunities and the corresponding median of the posterior distribution of OU’s α, half-life and *w* slope. *A, C, E*) Uniform OU’s α prior. *B, D, F*) Half-life derived OU’s α prior. Dashed lines delimit confidence intervals (95%) of predictions.

As the OU’s α prior mode was found to be the best predictor of the median posterior distribution of OU’s α, we built the causal models considering only the tests performed using each OU’s α prior mode (‘uniform’ or ‘half-life’). Figures 5*A*, *B* show models built considering the uniform OU’s α prior, while figures 5*C*, *D* show models built for the half-life OU’s α prior. In general, models using the half-life derived OU’s α had worse overall fit than models using the uniform OU’s α prior, which had lower Fisher’s C statistics than the former. Within the models using the uniform OU’s α prior, the most adequate scenario was obtained for the scenario considering log-transformed real half-life explaining the median posterior distribution of half-life, considering the uniform OU’s α prior (*p* = 0.67; fig. 5*B*). Furthermore, that model showed the lowest correlation between the median of the posterior distribution of OU’s α and *w* slope.

**Figure 5:**
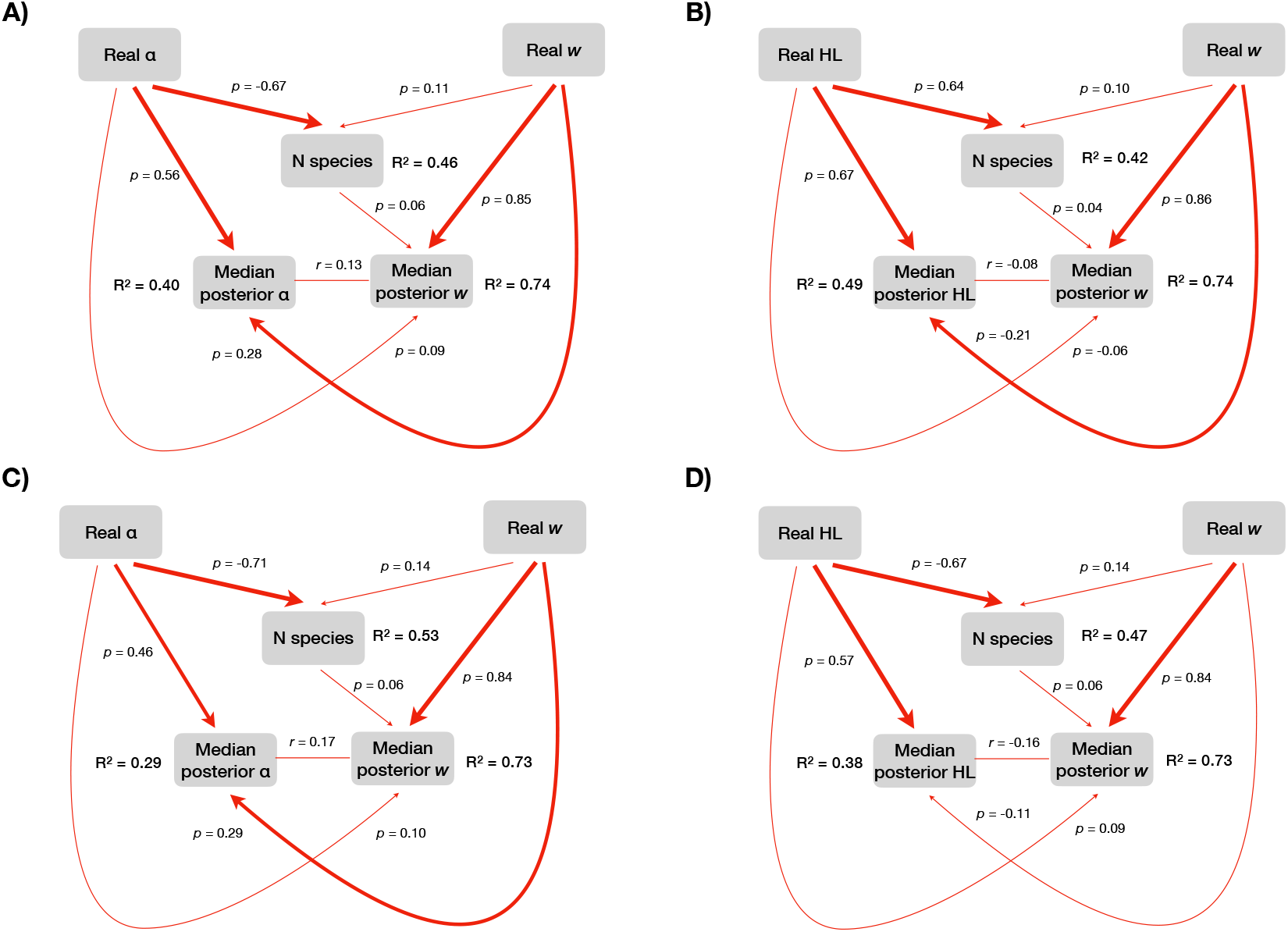
Causal models relating real OU’s α or half-life, real *w* slope and number of species in 1,000 simulated metacommunities on the median of the posterior distribution of OU’s α of half-life and *w* slope estimated using uniform (*A*, *B*) or half-life derived (*C, D*) OU’s α prior. Causal relationships were estimated using ordinary least squares. Half-life and *w* slopes were log-transformed prior to analysis. Validity of overall models was tested using d-separation tests (Shipley 2000). *A*) OU’s α model, uniform α prior (Fisher’s C = 1.59. *P*_df=2_ = 0.45). *B*) Half-life model, uniform α prior (Fisher’s C = 0.63, *P*_df=2_ = 0.73). *C*) OU’s α model, half-life α prior (Fisher’s C = 4.13, *P*_df=2_ = 0.13). *D*) Half-life model, half-life α prior (Fisher’s C = 4.64, *P*_df=2_ = 0.10). Arrow width indicates the strength of the respective causal relationship.

### Phyllostomid bats

ABC analysis of phyllostomid bats generated a prior distribution of OU’s α and *w* slope parameters containing 16,040 simulated metacommunities. Both prior and posterior distribution of parameters are shown in figure 6. The posterior distribution of OU’s α (fig. 6*A*) indicated a density peak at α = 0.05, which means a niche half-life (t_1/2_ =) 13.9 Myr, or 41% of the total timespan of the phylogenetic tree of phyllostomids (34 Myr). That is to say that the distribution of entropy across the temperature seasonality gradient showed high phylogenetic signal.

**Figure 6:**
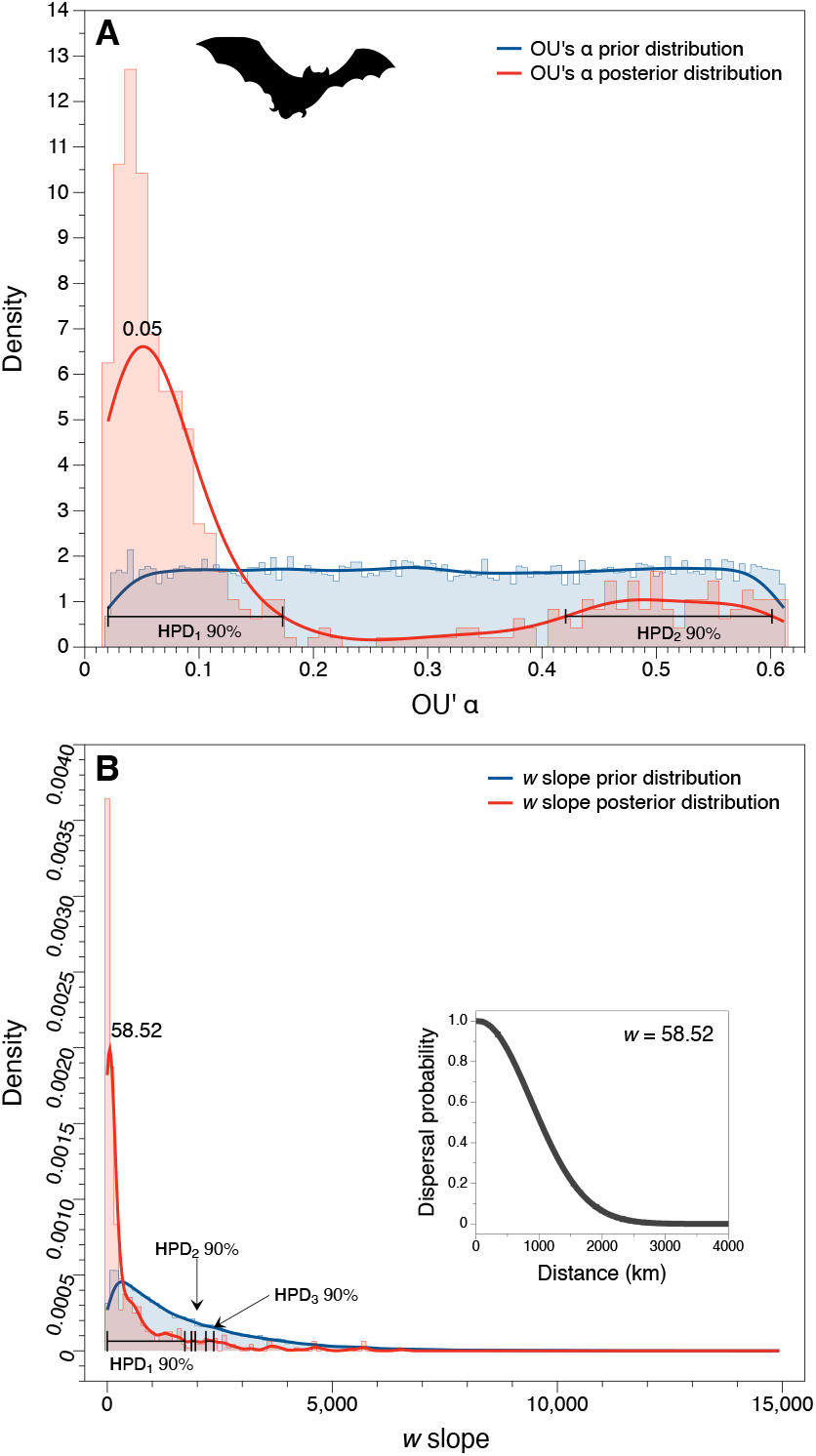
Histograms and density plots of the posterior distribution of (*A*) OU’s α and (*B*) *w* slope parameters underlying species diversity of phyllostomid bats along a gradient in temperature seasonality in the Neotropic. Parameters were estimated using ABC analysis implemented in *mcfly* package. Prior sample size: 16,040 simulations. Posterior sample size: 480 simulations. Only the most representative highest posterior density intervals (HPD_1_ 90%) were interpreted.

Moreover, the analysis estimated a peak density for *w* slope = 58.52. The minimum spanning tree computed based on the geographic coordinates of sites indicated that the maximum distance necessary to connect all cells in the map was 263 kilometers. Based on eq. 3, at such distance the probability of a species to disperse from a site to another was very high (*δ_lk_* > 0.96, see the subplot within fig. 6*B*); the probability δ_*lk*_ only decreases to 0.5 at distances ≡ 1,000 kilometers.

Figure 7*A* shows the observed entropy distribution of bat assemblages across the Neotropic, while figure 7*B* provides mean estimates of entropy computed from those simulations whose OU’s α and *w* slope parameters fell within the boundaries of the most representative HPD computed for both parameters. As we can see, ABC algorithm captured most entropy variation across sites. In figure 7*C*, we plotted the residuals of a cubic model (subplot within fig. 7*C*) relating the observed entropy of sites (y-axis) to ABC-estimated entropy (x-axis). As we can see, although there is a strong association between entropies, it is not quite linear, especially at lower and upper limits of entropy values. Interestingly, the residual plot indicates that in the southernmost regions of the Neotropic, where entropy values were lower, the simulation framework either under or overestimated the entropy of bat assemblages (fig. 7*C*).

**Figure 7:**
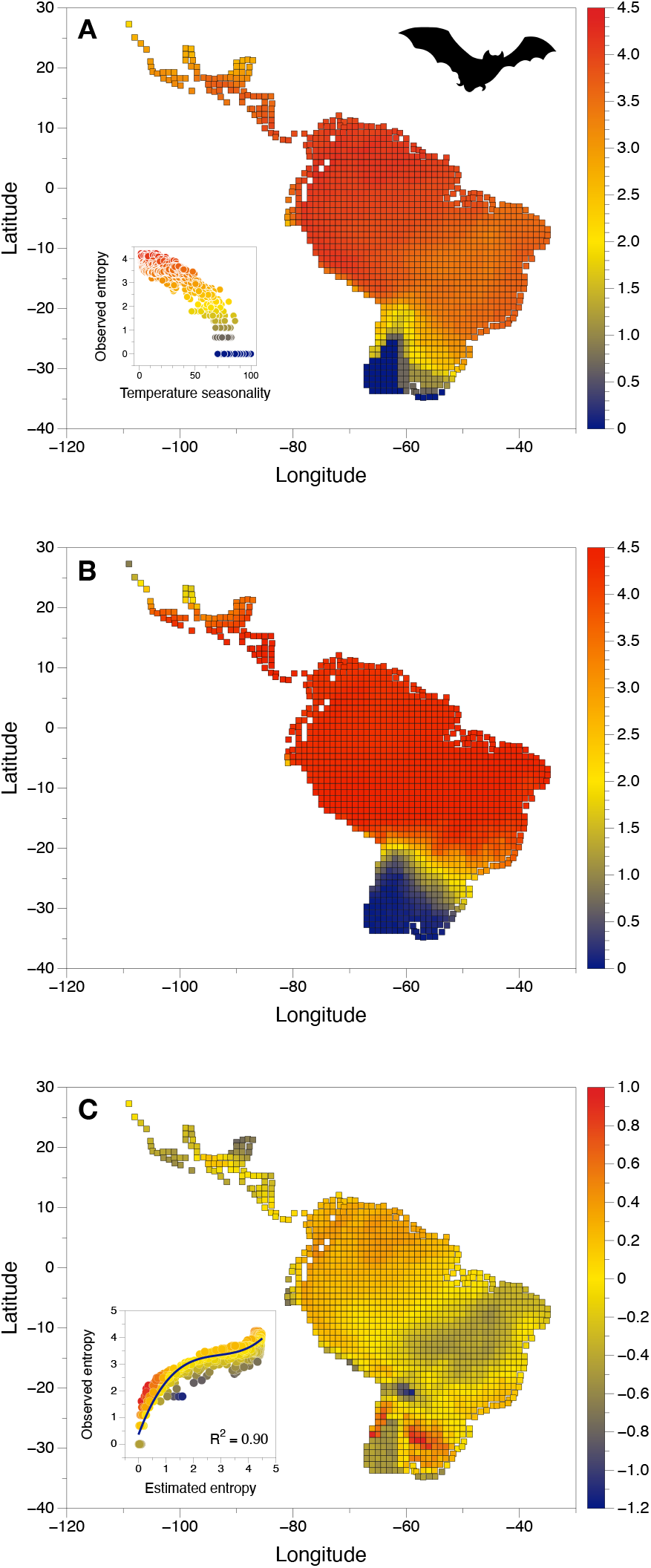
Species diversity gradients of phyllostomid bats across the Neotropic. (*A*) Observed entropy. The subplot within shows strong negative correlation between entropy and temperature seasonality, with highest entropy values associated with lower temperature seasonality. (*B*) mean entropy estimated using ABC analysis. Mean values were computed for simulations with OU’s α and *w* slopes falling within the boundaries of the most representative HPD interval (see fig. 6) computed for the corresponding posterior distribution of each estimated parameter. (*C*) Residual values of the cubic model relating estimated (x-axis) and observed entropy (y-axis) (see subplot within the main plot).

## Discussion

The framework proposed in this study and implemented in the package *mcfly* successfully enabled us to infer the most plausible scenario of niche evolution determining biological diversity gradients across spatially-structured environmental gradients. We demonstrated that our model-based simulation approach, coupled to rejection-based Approximate Bayesian Computation (ABC) algorithm (Beaumont 2010; Csilléry et al. 2010), allows to estimate (i) the strength of adaptation rate along the niche evolution of species from a monophyletic lineage and distributed across metacommunities; and (ii) the role played by dispersal limitation on the distribution of biological diversity across space. Considering the initial aims of community phylogenetics proposed by Webb et al. (2002), our approach provides meaningful estimates of the role played by phylogenetic relationships on species assembly patterns, and helps to reconcile niche and neutral processes, a topic heatedly debated over the last decades (Vellend 2016, and references therein). Previous studies have already pointed out the need of integrating niche evolution to diversity gradients (Cadotte et al. 2017). This has been explored with correlative approaches relating phylogenetic and trait diversity (Tucker et al. 2018) or, indirectly, by estimating phylogenetic signal in one or more traits used to compute functional diversity (Muscarella et al. 2016). However, we demonstrated it is possible to link niche evolution models and species diversity gradients without assuming a priori phenotypic traits to define species niches. This releases the need of assuming a given trait (or group of traits) as the most appropriate for assessing niche-environment associations (why not any other[s]?). An advantage of our approach is that it allows the evaluation of datasets for which a complete characterization of niche dimensions is currently not available. Thus, it may fill an important gap in the knowledge on the relationship between phylogenetic relationships among species and ecological diversity.

The analyses performed using simulated datasets demonstrated that the ABC approach implemented in *mcfly* is suitable to estimate the adaptation rate of species niche positions and the degree of dispersal limitation across the assemblages in a metacommunity based on species diversity variation across environmental gradients. Nonetheless, the analytical framework showed to be sensitive to the mode of the prior OU’s α distribution (Roos et al. 2015). Therefore, we recommend the use of the uniform distribution of OU’s α to avoid biased posterior distributions of OU’s α parameter. Causal models detected solid links between real and estimated niche parameters (and for the dispersal parameter - *w* slope), either where OU’s α or niche half-life was considered. Thus, even though a formally defined function relating biological diversity to evolutionary models of niche evolution is not available to date, the ABC approach implemented in the package *mcfly* proved to be a useful way to estimate the imprint of niche evolution on diversity gradients (Cadotte et al. 2017). Although evolutionary analyses usually suffer from incomplete datasets (FitzJohn et al. 2009; Nee et al. 1994; Pybus and Harvey 2000), our framework demonstrated to be robust even for metacommunities showing incomplete phylogenetic data (simulated metacommunities contained at least 40% of total phylogenetic pool), which is quite common in ecological and biogeographic data. Yet, some caution may be necessary where regional assembly contains only a small fraction of the species within the phylogenetic tree.

Phyllostomid bats showed strong dependency on niche evolution towards an optimum of low temperature seasonality across Neotropical assemblages, corroborating results obtained by Stevens (2011). Phyllostomids have high dispersal capacity up to distances of *ca.* 1000 km, which is in conformity with the large geographic range of most of its species (Willig et al. 2003). Moreover, the estimated OU’α of phyllostomid’s niche positions was low (α = 0.05), which confers a relatively high half-life to niche positions of species (13.9 Myr). Those results suggest that those bats preferably track habitats offering niche conditions close to their optimum over long distances, instead of quickly adapting their niches towards an optimum condition (see also Villalobos et al. 2013). That strategy was already hypothesized to explain plant distribution (Donoghue 2008). This finding may explain evidence showing that closely related phyllostomid species tend to live in geographic proximity (Villalobos et al. 2014).

In general, estimated entropy closely mirrored observed richness in bat assemblages, although not quite linearly, as revealed by the model relating observed to estimated entropy. For instance, at higher observed entropy levels in Amazonia region, simulations tended to slightly underestimate entropy. Certainly, another relevant niche factors might enable diversity levels to be higher than those expected by climatic niche alone. Vertical stratification in tropical rainforests, associated with seasonal floods affecting nutrient and roost availability, is a well-known driver of local phyllostomid diversity in Amazonian forests (Bernard 2001; Pereira et al. 2010; Pereira et al. 2009). Some other remarkable deviation patterns were observed in two regions: (1) Southern Atlantic forests, where entropy was strongly underestimated. Here, phyllostomids are a small subsample of the diversity of the family, as may be inferred from the IUCN distribution maps (Rojas et al. 2018a), and are mostly represented by generalist frugivores who present higher speciation rates than phyllostomids of other feeding guilds (Rojas et al. 2018b). Such generalist diet would allow them to cope with the unpredictability of food resources in the subtropical regions. (2) The “dry” diagonal of South America, where entropy was in general overestimated. Neotropical rainforests and savannas share a significant part of the generalist frugivore assemblage, but the absence of dense forests in the more open physiognomies negatively affects the occurrence of several frugivorous phyllostomids (Stevens 2006), particularly those specialized in canopy fruits. Therefore, the phyllostomid assemblages of the South American “dry” diagonal become more represented by obligatory, and specialized, nectarivores, which show lower rates of speciation than those observed for the generalist herbivore lineages (Rojas et al. 2018b); this might explain entropy overestimation. In all cases, a single optimum niche condition might fail to account for other relevant variables such as vegetation types. Indeed, the current version of *mcfly* has this limitation, which should be fixed in further developments of the analytical framework. Yet, in its current version, *mcfly* allows for multiple conditions to be included as a principal component, and users may thereby incorporate climatic and landscape data to the analysis. Finally, mismatches between observed and estimated entropy in some areas may be, to some extent, due to inconsistencies in IUCN maps.

The framework proposed here shed light on the link between niche evolution and the distribution of biological diversity, and thereby improved our understanding of evolutionary imprints on ecological patterns. Yet, we believe that the main strength of this new approach is to open avenues of investigation that may help solving the eco-evolutionary puzzle. For instance, multiple niche optima, possibly derived from different niche conditions, and affecting differentially distinct clades, might provide more realistic scenarios than the single optimum scenario developed here. Further, our framework deals with evolutionary imprints on ecological patterns, but it does not allow evaluating how higher ecological mechanisms (sensu Vellend 2016) may ultimately affect evolutionary dynamics (Weber et al. 2017). Further developments of the framework proposed here should help filling these (and other) gaps in near future. This tale is to be continued.

## Acknowledgements

LD, MVC, MJRP and JAFDF research activities have been supported by CNPq Productivity Fellowship (grants 307527/2018-2, 306590/2018-2, 311297/2018-8 and 301799/2016-4, respectively). LD, GN, MVC and JAFDF research have been conducted in the context of the National Institutes for Science and Technology (INCT) in Ecology, Evolution and Biodiversity Conservation, supported by MCTIC/CNPq (proc. 465610/2014-5) and FAPEG (proc. 201810267000023). The PhyloPic (http://phylopic.org) image of a phyllostomid bat used in figures 6 and 7 was created by Melissa Ingala, and its use was authorized under the license available at https://creativecommons.org/licenses/by/3.0/. Figure 1 was brilliantly designed by Clara Heinrich (linkedin.com/in/clara-heinrich-4bb3a3112).

A ciência brasileira resiste.

